# Deep learning-enabled phenotyping reveals distinct patterns of neurodegeneration induced by aging and cold-shock

**DOI:** 10.1101/2020.03.08.982074

**Authors:** Sahand Saberi-Bosari, Kevin B. Flores, Adriana San-Miguel

## Abstract

Access to quantitative information is crucial to obtain a deeper understanding of biological systems. In addition to being low-throughput, traditional image-based analysis is mostly limited to error-prone qualitative or semi-quantitative assessment of phenotypes, particularly for complex subcellular morphologies. In this work, we apply deep learning to perform quantitative image-based analysis of complex neurodegeneration patterns exhibited by the PVD neuron in *C. elegans*. We apply a Convolutional Neural Network algorithm (Mask R-CNN) to identify neurodegenerative sub-cellular protrusions that appear after cold-shock or as a result of aging. A multiparametric phenotypic profile captures the unique morphological changes induced by each perturbation. We identify that acute cold-shock-induced neurodegeneration is reversible and depends on rearing temperature, and importantly, that aging and cold-shock induce distinct neuronal beading patterns.

## 1. Introduction

Aging, environmental stressors, and injury can induce reversible or irreversible changes at the subcellular, cellular, and tissue levels of an organism^1–11^. The *Caenorhabditis elegans* nervous system is not an exception and undergoes morphological and functional deterioration under these conditions. Morphological phenotypes indicative of neurodegeneration in this roundworm include somatic outgrowth, distorted soma, branched and wavy dendrites, and dendritic beading^2,8,12–18^. The ability of neurons to recover from degeneration has also been studied. For instance, Oren-Suissa et al. found that primary dendrites in the PVD neuron reconnect via branch fusion following laser surgery^19^. PVD is a widely studied multi-dendritic nociceptor neuron that responds to harsh touch (mechanosensor) and cold temperatures (thermosensor) (**Figure 1a**)^20–27^. Prior work has identified genetic pathways important for organization of dendritic branches and dendritic self-avoidance^28–33^. Dendritic organization in PVD is also affected by aging; while young animals have well-organized menorah-like dendritic structures, these tend to be replaced by non-uniform and chaotic outgrowth of dendritic branches^32^. Recently, Lezi et al. identified the formation of protrusions (or beading) along the dendrites of PVD during aging, through a process driven by the expression of an antimicrobial peptide^34^.

**Figure 1.**
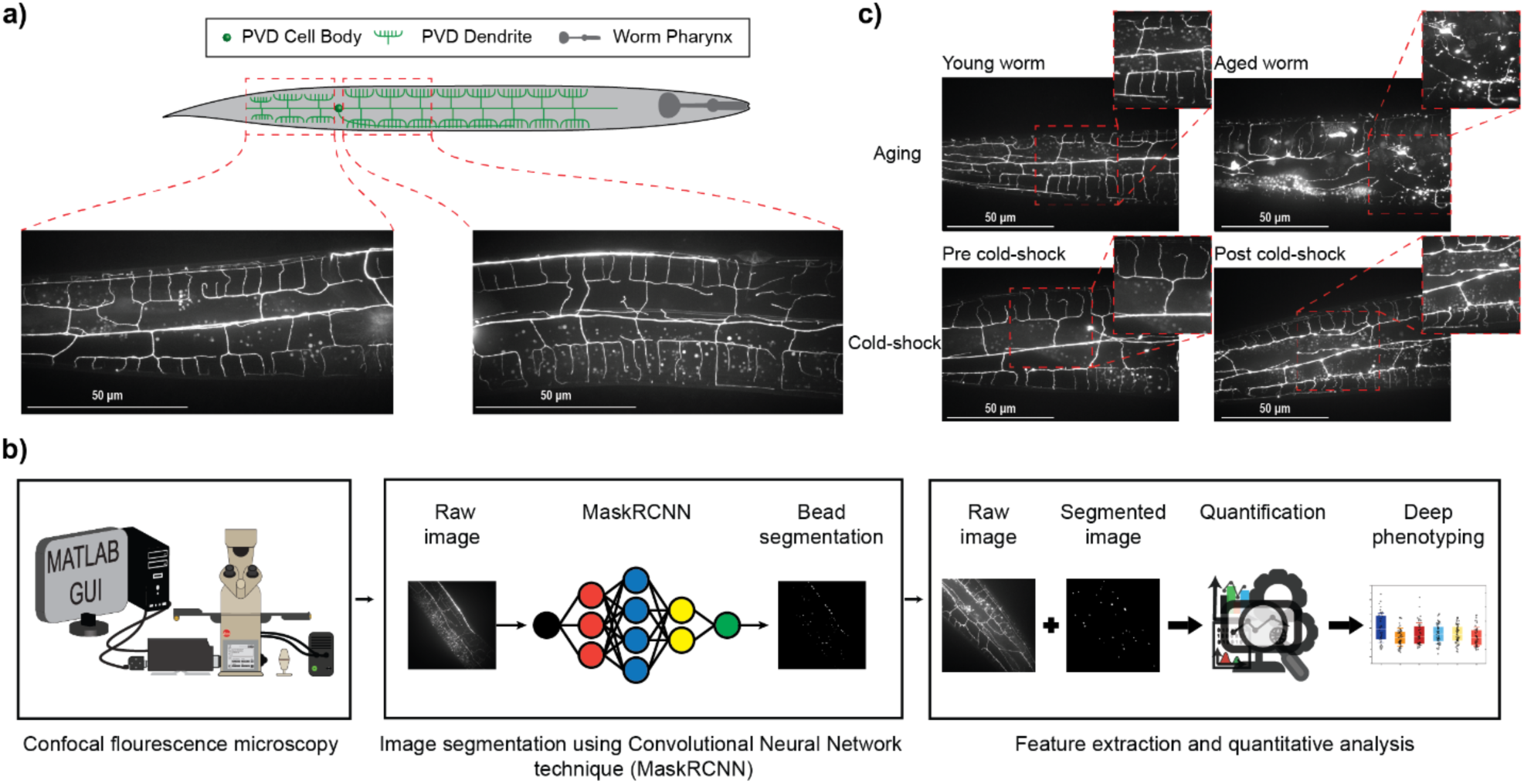
Quantitative analysis of PVD neurodegeneration by deep learning. **a)** Schematic of PVD neuron with menorah-like dendritic branches. Fluorescence images of PVD anterior and posterior to the cell body. **b)** Schematic of quantitative analysis pipeline to study PVD neurodegeneration. **c)** Aging and acute cold-shock induce neurodegeneration on PVD dendrites. These two stressors increase the formation of bubble-like protrusions along the dendritic arbors of PVD.

Characterization of PVD beading has thus far been performed by visual inspection and manual counting of fluorescent images, which is labor-intensive, time-consuming, and does not provide additional information about the observed morphological changes, aside from number of beads. Traditional image processing approaches typically rely on intensity difference for image segmentation^35–37^. The protrusions that appear in PVD have fluorescence intensities similar to the rest of neuron and autofluorescent lipid droplets. Thus, traditional image processing approaches are unable to perform the challenging segmentation of PVD protrusions. Quantitative analysis of PVD neurodegeneration morphology is important to understand the root causes of neurodegeneration. Machine learning has proven useful for analysis of biological systems and deep phenotyping^9,38–40^. In this work, we sought to integrate cutting-edge deep learning approaches to segment beads in PVD fluorescence images from live animals (**Figure 1b**). Convolutional Neural Networks (CNNs) have recently shown state-of-the-art performance in image segmentation tasks across a wide range of biological and biomedical images datasets^41–47^. Here, we utilize Mask R-CNN^48^, a CNN model that is designed to predict binary instance masks (one mask per predicted bead object) from an image to detect PVD beads. We follow this user-free segmentation approach with multiparametric phenotyping of PVD by extracting 46 quantitative features that describe beading patterns. These metrics include number of beads, cumulative area occupied by beads, average bead size, average pair-wise inter-bead distance, etc. We take advantage of the quantitative data provided by this pipeline to track subtle neurodegenerative phenotypes caused by different physiological stressors (**Figure 1c**). We validate our pipeline by assessing the effects of aging on PVD beading, and recapitulate previously observed changes^34^. In addition, we identify a previously unknown degenerative effect of exposure to acute cold-shock on neuronal structure. Finally, we show that this deep phenotyping approach enables predicting the biological status of a nematode (young, aged, cold-shocked) based on the quantitative metrics generated by the pipeline with over 85% accuracy. This analysis reveals that different stressors (aging and cold-shock) induce distinct neurodegenerative phenotypes hinting at potentially different underlying neurodegeneration mechanisms. This approach enabled automating image analysis of PVD neurodegeneration thus increasing throughput, eliminating the human bias and error introduced by manual assessment, and facilitated high content quantification of the subtle neurodegenerative changes in PVD, unfeasible in conventional methods.

## 2. Results and Discussion

### 2.1 Training the Mask R-CNN algorithm to perform complex image segmentation

We adapted the convolutional neural network (CNN) model Mask R-CNN^48^ to automatically detect bead protrusions in high resolution images of nematode dendrites (**Figure 2a**). The input to Mask R-CNN is a 1-channel grayscale microscopy image (1024 × 1024 × 1) and the output is a set of predicted bead regions consisting of one binary instance mask (1024 × 1024 × 1) per bead, i.e., a pixel has a value of 1 in the mask when it is part of a bead and 0 otherwise. A tiling procedure was employed to adapt Mask R-CNN for use with 2048 × 2048 × 1 microscopy images (see Methods and Materials), since this image size was sufficient to resolve the smallest bead protrusions. The Mask R-CNN architecture first generates regions of interest (ROIs) using a Faster R-CNN model, composed of a residual network (ResNet-101^49^) and a feature pyramid network^50^. ROIs are then processed with region proposal and ROI align neural network layers to produce an instance segmentation mask for each detected object. In contrast to thresholding based methods, which only rely on image intensity for predicting segmentations, CNNs automatically learn and then use hierarchical sets of image features directly from the training data without requiring manual feature engineering. Learning features enables relevant local context to be used in making segmentation predictions, e.g., the shape and size of the bead, what a dendrite looks like, and the proximity of beads to dendrites. We leveraged a transfer learning^51^ approach in which Mask R-CNN is pre-trained on a large annotated dataset (ImageNet^52^), and then fine tuned on a data set of nematode images that we manually annotated.

**Figure 2.**
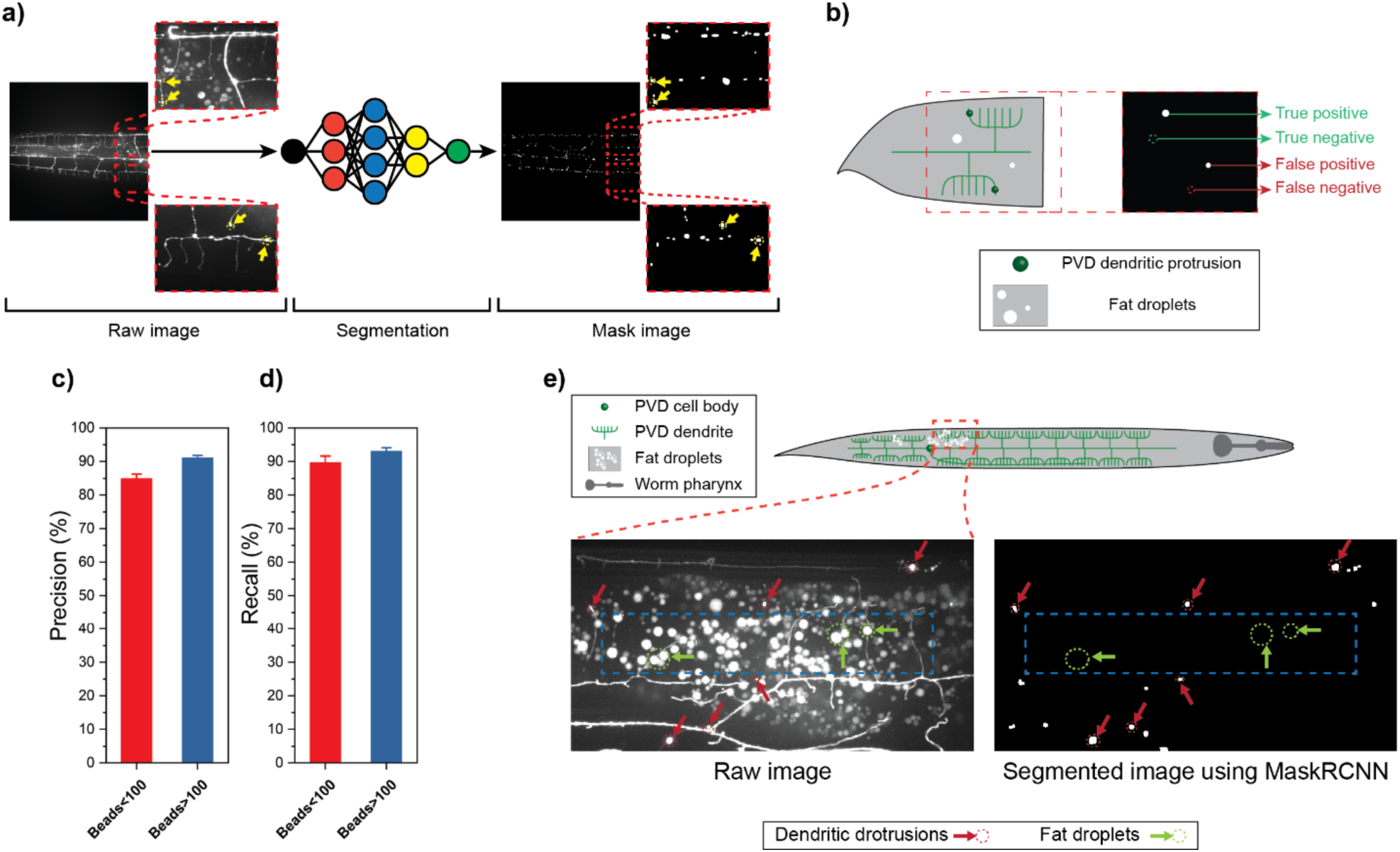
Deep learning approach successfully identifies beads in the PVD neuron. **a)** Schematic of segmentation pipeline. Raw 2048×2048 images are fed to the trained Mask R-CNN model to perform instance segmentation. Yellow arrows point to neuronal beads. **b)** Illustration for true positive, true negative, false positive, and false negative cases used to quantify the performance of instance segmentation. **c-d)** The performance of the algorithm was examined by defining precision and recall of segmentation where 12 validation images were used. Error bars are Standard Error of Mean (SEM). **e)** Images showing the algorithm successfully distinguishes bubble-like protrusions (beads) from fat droplets.

The Mask R-CNN algorithm requires a training data set comprised of raw images of PVD and their corresponding ground truth masks that label the protrusions. The masks were created from raw images using a custom MATLAB code that allows the user to draw around each bead location. A total of 19 images (each with ∼ 50-150 beads with an average size of ∼150 pixels) were manually segmented to compile the training set. In addition, an independent validation set was generated with 12 raw images and their associated binary masks. The validation set includes diverse images with ∼30 to ∼150 beads. These were equally split into images with a low (<100) and a high (≥ 100) number of beads, to test segmentation consistency. To assess segmentation performance, we quantified precision and recall, described as:

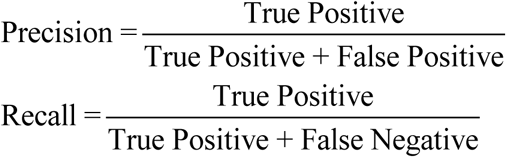

In these expressions, true positives are correctly identified beads, false positives are non-bead objects identified as beads, and false negatives are non-identified beads (**Figure 2b**). As shown in **Figure 2c**, the segmentation precision for the validation data set was 85% and 91% for images with low and high bead numbers, respectively. Similarly, a recall of 90% and 93% was obtained for low and high bead number images, respectively (**Figure 2d**). These slight differences could stem from the low number of beads while retaining the same level of objects that can be falsely identified as beads in the first group The optimized Mask R-CNN algorithm successfully scored 88% in precision and 91% in recall for the entire validation set. Thus, this machine-learning approach offers consistent unbiased segmentation with high accuracy.

Importantly, precision and recall do not provide information to assess the performance of the model in ignoring objects that can easily be identified as beads (true negatives). In this particular phenotyping problem, this type of objects are prevalent. Autofluorescent lipid droplets can be easily mistaken for neurite protrusions, due to their round shape and location, which can overlap with PVD dendrites in maximum projections. Distinguishing round objects with comparable intensity levels and with similar locations and sizes is a significant challenge. To assess the power of the algorithm to distinguish between the two, we chose 3 images from animals with an abundance of fat-droplets that overlapped with dendrites, as part of our training set. As shown in **Figure 2e**, the algorithm is successful in discerning fat droplets from beads, despite their similarities. Prior approaches have addressed this problem by performing dual color microscopy to compare images that show only lipid droplets with images that show the fluorescent reporter^38^. This deep learning approach eliminates the need to perform alternative analyses or dual color microscopy to subtract autofluorescent objects.

### 2.2 Deep phenotyping of age induced PVD neurodegeneration

The nervous system in *C. elegans* undergoes morphological and functional decline due to aging^14,18^. Morphological changes in PVD include dendritic outgrowth and beading, which become more common as animals age, as evidenced in **Figure 3a**. As previously mentioned, quantitatively investigating beading is difficult as animals can exhibit tens to hundreds of beads with fluorescence intensity levels similar to those of labeled neurons and autofluorescent lipid droplets. Moreover, beading is a highly variable process, and quantification thus requires analysis of large animal populations. We first aimed to quantitatively analyze aging-induced beading in PVD using the deep learning pipeline. Our results (**Figure 3b**) show that the average bead count increases from days 2 to 4 and 6 of adulthood. Interestingly, the average number of protrusions does not appear to change significantly afterwards. These results suggest that there may be a saturation point for the beading process, which animals reach at mid-age.

**Figure 3.**
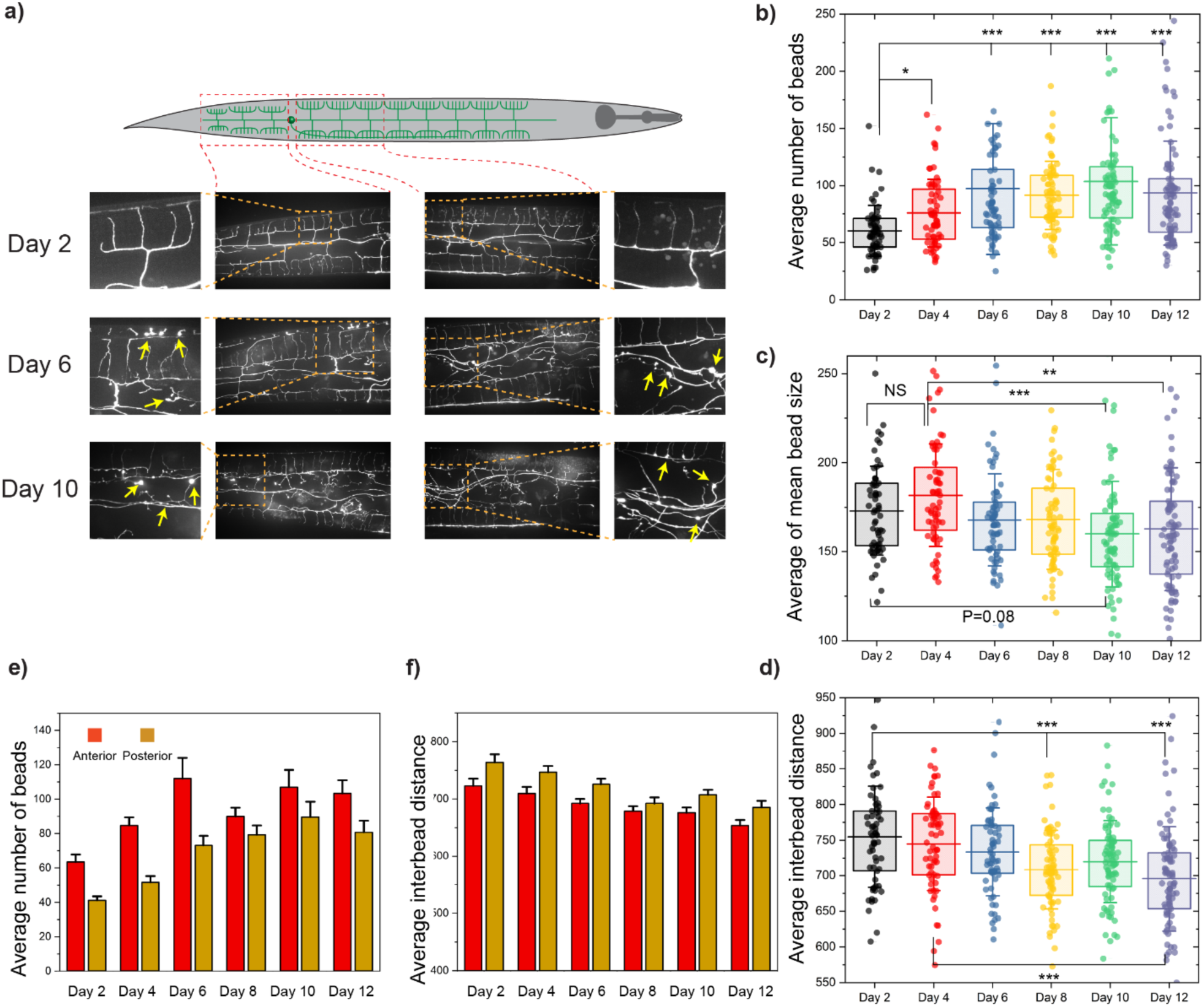
Deep learning allows quantitative analysis of aging induced morphological changes in PVD. **a)** Qualitative inspection of PVD at 3 time points of their life-span shows an increase in number of beads throughout the dendrites. Protrusion formation was identified in both anterior and posterior parts of the PVD neuron. Yellow arrows point to neuronal beads. **b-d)** Average number of beads, average of mean bead size, and average inter-bead distance of both anterior and posterior regions of PVD throughout aging. Lines are 25^th^ percentile, mean, and 75^th^ percentile. Whisker is standard deviation. Statistical analysis was performed with one-way ANOVA followed by Tukey multiple comparison correction. *P<0.05, **P<0.001, and ***P<0.0001. **e-f)** Average number of beads, and average inter-bead distance of anterior versus posterior regions of PVD throughout the aging process. Error bar is SEM.

One of the advantages of computer-based image segmentation is that quantification of beading neurodegeneration is not limited to the number of beads. Our post-segmentation MATLAB pipeline enabled extracting additional metrics (a total of 46, Supplemental information) to comprehensively describe the morphological neurodegeneration phenotypes. The average bead size (**Figure 3c**) seems to decrease slightly as animals age (days 6-12 vs. days 2-4), which can be explained by an increase in percentage of small beads (area < 100 pixels) (**Figure S1b**). While the size is slightly reduced, the total area occupied by beads increases as nematodes age (**Figure S1a**). These results suggest that the main morphological change induced by aging is an increase in total beading (as measured by number or total bead area), rather than in bead size. The average inter-bead distance (i.e., average of all pairwise distances) describes how dispersed the beads are, and decreases in older populations as expected due to an increase in total number of beads (**Figure 3d**). Other metrics that describe bead size and spatial bead distribution (such as 90^th^ percentile of bead size, and percentage of pairwise inter-bead distances < 300 pixels, **Figures S1d-e**) confirmed an overall trend towards accumulation of smaller beads with increased density throughout the neuron in older animals.

To deepen our understanding of aging-induced beading, we compared the patterns exhibited anterior (towards the head) and posterior (towards the tail) to the PVD cell body, since separate images were acquired (**Figure 3a**). While both regions exhibit an increase in number of beads (**Figure 3e**), this change was more drastic in the anterior section. This difference could be explained by either a higher susceptibility to beading or by the fact that the anterior region occupies larger area, since the posterior is closer to the animal’s tail and is thus more tapered. The average inter-bead distance in the posterior region tends to be larger than in the anterior side (**Figure 3f**), as would be expected for a reduced number of beads. As shown in **Figure S1f**, bead morphology appears to be homogeneous, as there is no significant difference in anterior vs. posterior average bead size. Metrics such as the percentage of small beads (< 100 pixels) or the percentage of beads with close neighbors (pairwise inter-bead distances <300 pixels) did not show any significant differences along the two different sections of PVD (**Figure S1g-h**). This deep learning – based analysis corroborates the neuronal beading reported by Lezi et al., while deepening our understanding of the subtle neurodegenerative patterns that result from aging.

### 2.3 Acute cold-shock induces neurodegeneration in PVD neuron

In addition to sensing harsh touch, PVD acts as a thermosensor activated by cold temperatures^53^. Cold-shock has been previously studied as a stressor for *C. elegans* ^53–64^. Robinson et al.identified that animals can survive short (4 hrs.) exposures to acute cold-chock (2 °C), but longer exposures (24 hrs.) result in death for a fraction of the population^65^. Furthermore, Ohta et al. showed that the pre cold-shock culture temperature is inversely correlated with survival rate (more animals survive cold-shock if previously cultured at lower temperatures)^66^. While the detrimental effects of cold-shock on nematodes’ survival and PVD’s involvement in responding to cold temperatures have been independently studied, the impact of cold-shock exposure on PVD health has not been investigated. To answer this question, we first tested the effects of exposure to cold-shock on PVD morphology, where we identified the appearance of PVD neurite beading. Thus, we sought to examine the neurodegenerative effects of acute cold-shock at 4 °C through our deep learning phenotyping pipeline.

To characterize the relation between cold-shock and beading, we first exposed different *C. elegans* populations to cold-shock for various durations. As shown in **Figure 4a**, eggs extracted from gravid hermaphrodites were transferred to NGM plates and cultured at 20 °C until day 2 of adulthood, when pre-cold-shock microscopy was performed. Nematodes were then split into four separate plates and transferred to 4 °C for either 4, 8, 16, or 24 hrs. Visual inspection of raw images suggested beading increases with longer cold-shock, but is especially evident in populations that were exposed for 16 hrs. or more. Quantitative analysis performed using the trained Mask R-CNN and post segmentation feature extraction pipeline shows that the number of beads gradually increases with longer periods of cold-shock (**Figure 4b**), and is almost doubled after 16 hrs., as compared to non-exposed animals. Similar to the aging process, beading reaches a saturation point, where no significant change in the number of beads is observed after 16 hrs. Interestingly, the percentage of small beads (area < 100 pixels) increases after 4 and 8 hrs. of cold-shock, but this effect is not observed after 16 and 24 hrs. (**Figure S2b**). This suggests that new small beads are generated in the first 8 hours, resulting in a higher percentage of smaller beads. The drop in percentage of small beads after 16 and 24 hrs. could be due to existing protrusions becoming larger once the number of beads saturate. This fluctuation in percentage of small beads is also reflected in the average size (**Figure 4c**), which slightly decreases during the first 8 hours of cold-shock and grows after 16 and 24 hrs. One potential explanation for these observations is that initially new small beads form, but eventually the beading mechanism switches to bead growth rather than bead generation.

**Figure 4.**
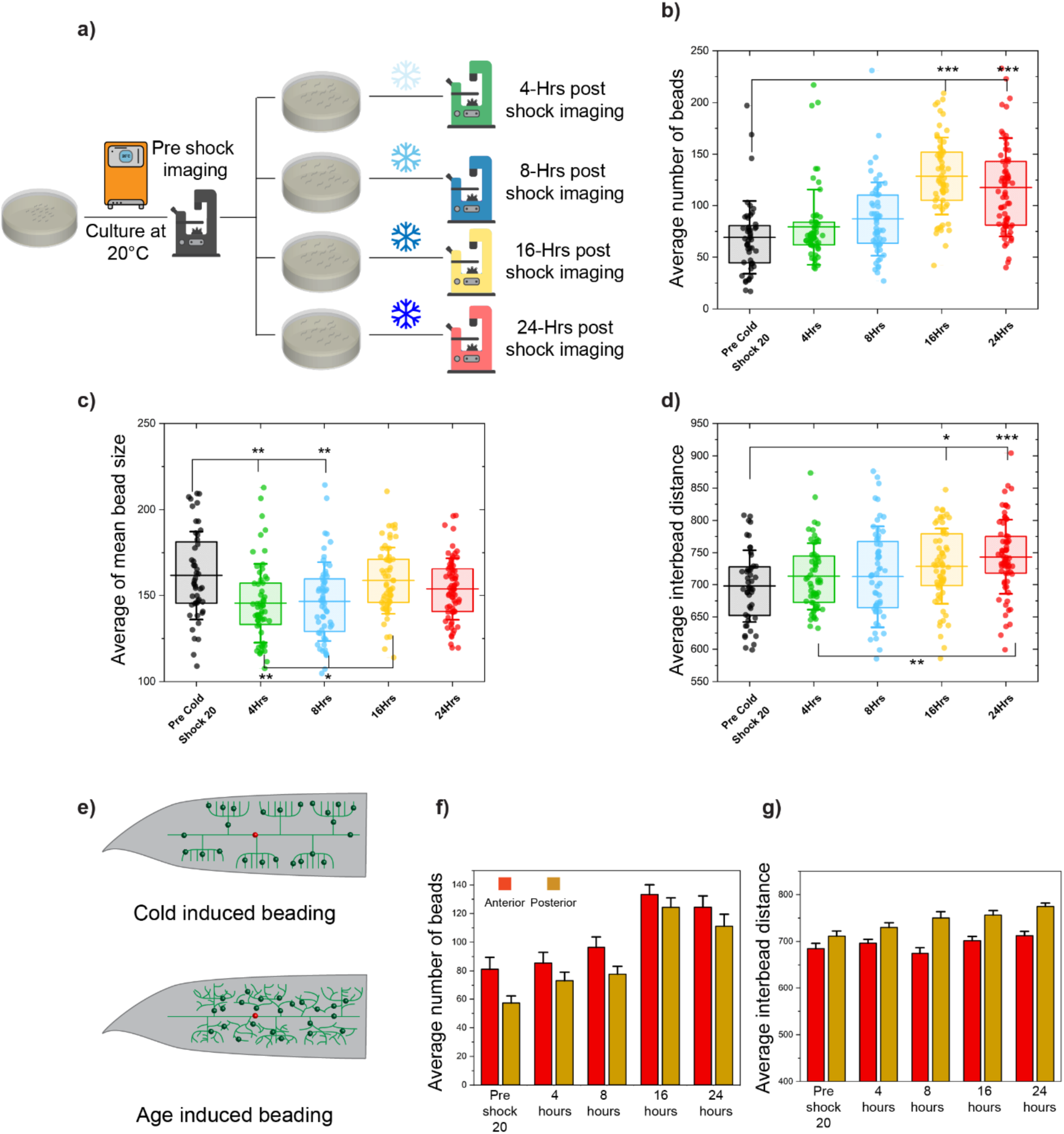
PVD neuronal structure undergoes morphological changes upon exposure to acute cold-shock. **a)** Schematic of acute cold-shock assay. Nematodes were cultured at 20°C until day two of adulthood, split into four plates, and cold-shocked for 4,8,16, or 24 hours. Fluorescence microscopy was conducted before exposure to cold-shock and after specific periods of shock. **b-d)** Average number of beads, average of mean bead size, and average inter-bead distance of anterior and posterior regions of PVD as nematodes experienced various duration of cold-shock. The lines are 25^th^ percentile, mean, and 75^th^ percentile. Whisker is standard deviation. Statistical analysis was performed with one-way ANOVA followed by Tukey multiple comparison correction. *P<0.05, **P<0.001, and ***P<0.0001. **e)** Illustration of distinct beading patterns in aging and acute cold-shock based on the inter-bead distance. Inter-bead distance decreases with aging while it increases with cold-shock. **f-g)** Average number of beads and average inter-bead distance of anterior versus posterior parts of PVD upon cold-shock. Error bar is SEM.

Computer-based image processing and quantitative analysis also enabled identifying subtle differences between aging and cold-shock beading patterns. While an increase in bead number was observed in both cases, cold-shock resulted in an increase in average inter-bead distance (**Figure 4d**), in contrast to aging. This counterintuitive result can potentially be explained by the tendency of cold-induced protrusions to form in more distant dendrites (such as 3^rd^ or 4^th^ order branches) of healthy menorahs. With aging, beads are generated evenly throughout the entire neuron, likely as a result of the aging-induced disorganized branching that increases the density of dendrites (where beads are formed) throughout the worm’s body (**Figure 4e**). The information extracted from anterior and posterior regions of PVD for nematodes exposed to acute cold-shock shows very similar patterns to aging-induced neurodegeneration. The number of beads in the anterior part is greater than in the posterior side (**Figure 4f**) and the beads are on average farther apart in areas closer to the tail (**Figure 4g**). Bead size appears to be homogeneous in both sides (**Figure S2f**), while the anterior region has a slightly higher percentage of small beads (area < 100 pixels) (**Figure S2g**). Utilizing this deep learning quantitative phenotyping enabled the identification of a previously unknown effect of acute cold-shock on PVD degeneration, which is exacerbated with longer exposures. Moreover, this analysis suggests that beading patterns differ for aging and acute cold-shock, suggesting potentially different mechanisms of protrusion formation.

### 2.4 Post cold-shock recovery can eliminate PVD dendritic protrusions

Given the significant increase in number of dendritic protrusions in PVD upon exposure to acute cold-shock, we next sought to determine its potential for regeneration. To test this hypothesis, we designed experiments to characterize PVD beading patterns after acute cold exposure and following a subsequent period under normal culture conditions (referred to as rehabilitation or recovery). As shown in **Figure 5a**, we performed 3 one-day rehabilitation regimes at 3 different temperatures, selected to cover the entire physiological range (15, 20, and 25 °C). Given that nematodes growth rate and life-span depend on culture temperature, we expected the population cultured at 25 °C to show a faster recovery rate than those grown at 15 °C. After exposure to 16 hrs. of acute cold-shock, the average number of beads increased by 100% as compared to pre cold-shock conditions. After one day of rehabilitation we observed a decrease in the number of dendritic protrusions in all three rehabilitation temperatures (**Figure 5b**). As expected, populations cultured at 15 °C and 25 °C had the lowest (∼30%) and highest (∼50%) recovery, respectively, suggesting that recovery rate is correlated with growth rate.

**Figure 5.**
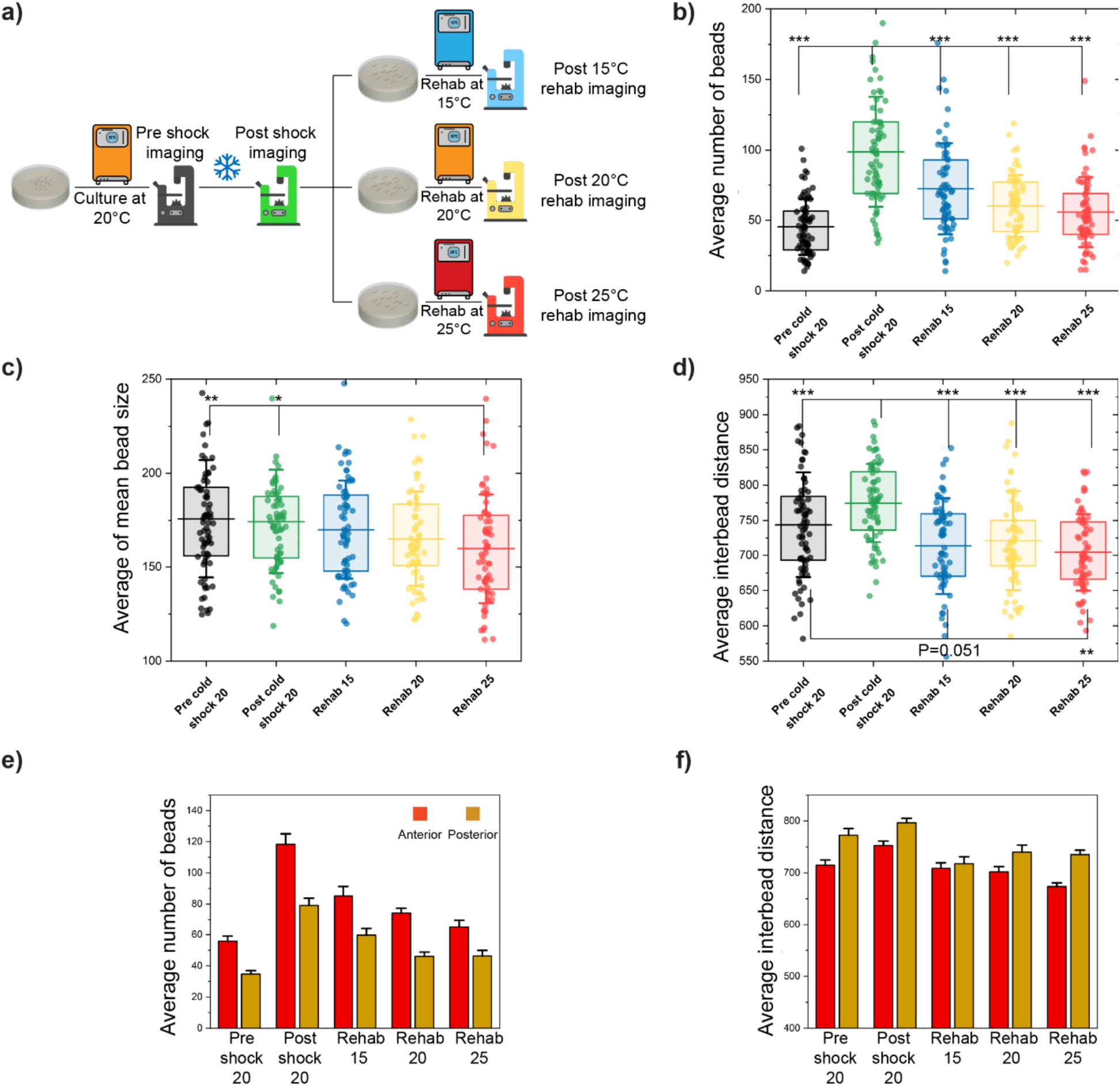
PVD neurodegenerative phenotypes caused by acute cold-shock are reversible. **a)** Schematic of post cold-shock rehabilitation treatment assay. Nematodes were cultured at 20 °C until day 2 of adulthood and were exposed to cold-shock for 16 hrs. To perform recovery, the population was split into three plates at either 15 °C, 20 °C, or 25 °C for one day. **b-d)** Average number of beads, average of mean bead size, and average inter-bead distance of PVD neuron as nematodes experienced cold-shock for 16 hrs. and undergo rehabilitation at 3 different temperatures. The lines are 25^th^ percentile, mean, and 75^th^ percentile. Whisker is standard deviation. Statistical analysis was performed with one-way ANOVA followed by Tukey multiple comparison correction. *P<0.05, **P<0.001, and ***P<0.0001. **e-f)** Average number of beads, and average inter-bead distance of anterior versus posterior regions of PVD as nematodes experienced cold-shock for 16 hrs. and rehabilitation at 3 different temperatures. Error bar is SEM.

In addition to a reduction in number, the average bead size slightly decreases after rehabilitation (**Figure 5c** and **Figure S4**). This recovery is corroborated by the total area covered by beads (**Figure S3a**), which increases after cold-shock and decreases in all recovery regimes, indicating that bead formation due to cold-shock is reversible. These results suggest that recovery occurs by both bead elimination and a gradual size reduction. To further understand the spatial patterns of cold-shock bead formation, we also explored inter-bead distances. As previously mentioned (**Figure 4c**), the average inter-bead distance increased post cold-shock, suggesting beads are formed in the farthest dendrites. One-day recovery treatment at all three temperatures reduced this metric (**Figure 5d**), suggesting that beads on the farthest dendrites are more prone to disappear post-recovery. As expected, the percentage of beads with close neighbors (inter-bead distance < 300 pixels) decreases with cold-shock and increases after recovery (**Figure S3d**).Taken together, these quantitative features suggest that cold-shock induces the formation of beads, particularly in distal regions (as the inter-bead distance increases), and that subsequent culture at physiological temperatures reverts these changes.

In line with previous findings, the anterior region of PVD exhibits a higher number of beads than the posterior region, post-rehabilitation. However, recovery does not appear to favor either side, as both areas show a reduction of beading post recovery (**Figure 5e**). Likewise, while the posterior region shows higher inter-bead distances than the anterior region, both exhibit a reduction of inter-bead distance post recovery (**Figure 5f**). The average bead size, percentage of small beads (area < 100 pixels), and percentage of beads with close neighbors (inter-bead distance < 300 pixels) do not show any significant differences between the anterior and posterior regions, either post cold-shock or post recovery (**Figure S3f-h**), for most conditions. This suggests that the propensity of the anterior region to increased beading observed with aging is also observed upon cold-shock and after recovery from cold-shock. Taken together, these results indicate that after acute cold exposure, one day recovery at different temperatures can almost completely alleviate the induced neurodegenerative effects. In addition, this data suggests that a more efficient recovery can be achieved by rehabilitation at higher temperatures. Finally, it appears that cold-shock preferentially induces beading in the farthest dendrites, but these are also preferentially removed during recovery.

### 2.5 Pre cold-shock culture temperature affects neurodegeneration severity

Physiological culture temperature is a key environmental factor that affects development, growth, and life-span in poikilotherms, such as *C. elegans*^57^. Nematodes habituate to imposed environmental conditions, including temperature^66–70^. Previous studies have identified that after a 4 °C of cold-shock over 85% of animals cultured at 25 °C die, while most animals cultured at 15 °C survive^66^. These findings motivated us to investigate whether the pre cold-shock culture temperature plays a role in beading neurodegeneration. To test this, 3 parallel cold-shock/recovery experiments at 3 physiological temperatures were conducted (**Figure 6a**) where populations were cultured at 15, 20, and 25 °C for ∼3.5, 2.5, and 1.5 days, respectively. These animals were then exposed to acute cold-shock at 4 °C and subsequently returned for one day to their culture temperature. The difference in culture time prior to cold-shock allowed animals to reach the same developmental stage. Based on prior studies where nematodes cultured at lower temperatures prior to cold-shock have a higher survival rate^66^, we hypothesized that lower temperatures would result in less severe neurodegeneration than high temperatures.

**Figure 6.**
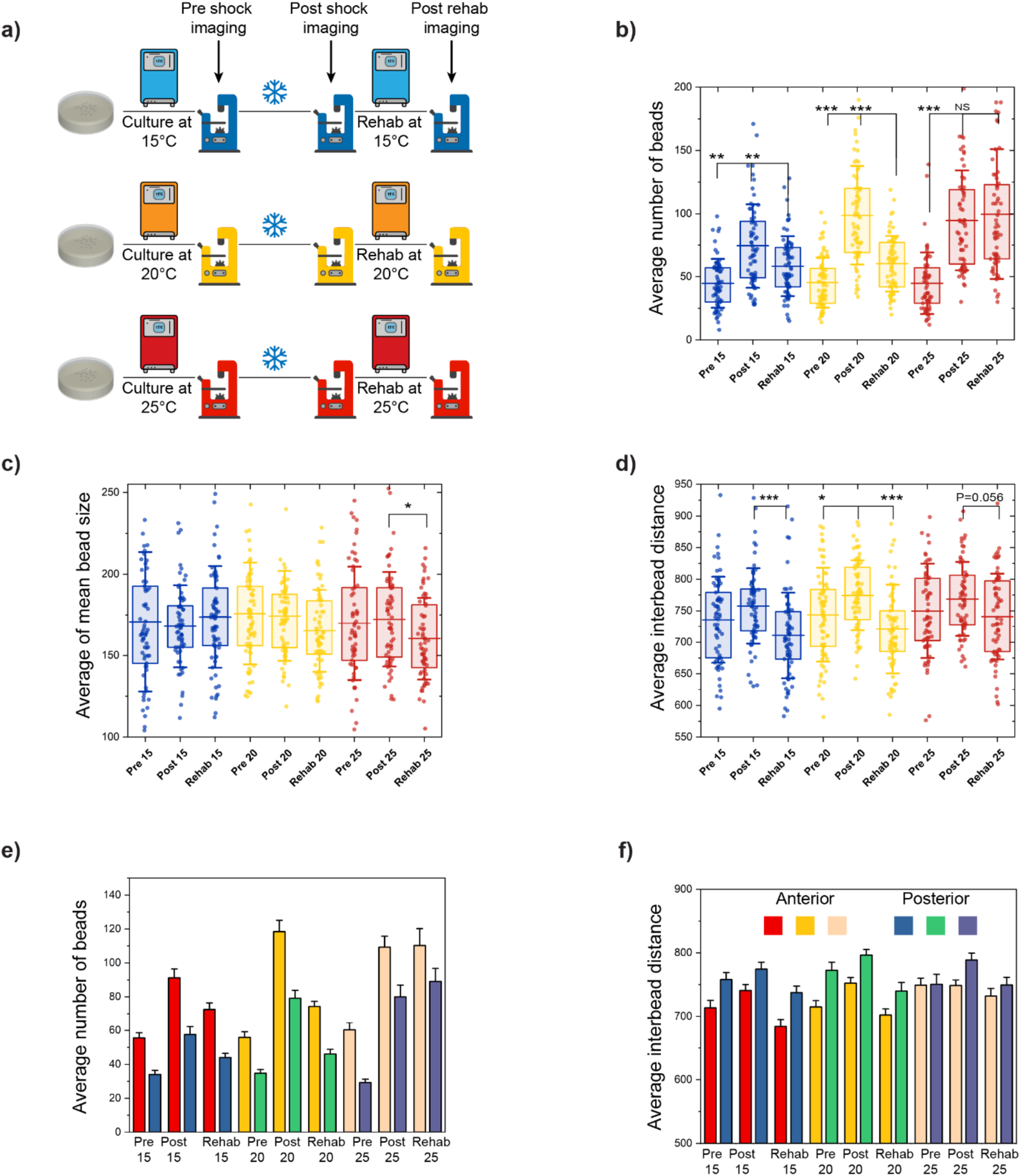
Populations cultured at different temperature before being exposed to cold-shock show different susceptibility to neurodegeneration. **a)** Schematic of the experimental setup to study the effect of pre cold-shock cultivation temperature. Animals were cultured at 20 °C until young adulthood and then transferred to 15 °C, 20 °C, or 25 °C for 3.5, 2.5, and 1.5 days, respectively. Cold-shock was then performed for 16 hrs. and rehabilitation was performed for one day at the pre cold-shock temperature. **b-d)** Average number of beads, average of mean bead size, and average inter-bead distance of PVD for populations that undergo cold-shock as described in part a). The lines are 25^th^ percentile, mean, and 75^th^ percentile. Whisker is standard deviation. Statistical analysis was performed with one-way ANOVA followed by Tukey multiple comparison correction. *P<0.05, **P<0.001, and ***P<0.0001. **e-f)** Average number of beads, and average inter-bead distance of anterior versus posterior part of PVD for populations that undergo cold-shock as described in part a). Error bar is SEM.

Post cold-shock behavioral analysis revealed that animals grown at 25 °C were the most affected, as they recovered mobility long after transfer to room temperature (30 – 40 min), while this time was considerably shorter for animals cultured at 15 and 20°C. Once animals started crawling, nematodes cultured at 25 °C moved significantly slower than those cultured at lower temperatures, indicating that worms habituated to a higher temperature undergo a more drastic shock under cold exposure. These observations suggest that a larger temperature gradient between culture and cold-shock results in increased neuronal damage. As shown in **Figure 6b**, the number of beads present after cold-shock and rehabilitation confirm this trend. The average number of beads increases post cold-shock in all samples, with the smallest change for nematodes grown at 15 °C. The mean bead count after cold-shock reaches the same level for samples cultured at 20 and 25 °C, potentially due to beading reaching a saturation point. This upper limit in number of beads was also observed in neurodegeneration caused by aging and in cold-shock exposure for different periods of time. Interestingly, while populations rehabilitated at 15 and 20 °C show a reduction in number of beads, this effect was not present in those recovered at 25 °C. This could be explained by either a delayed or slower regeneration, or an inability to regenerate for animals cultured at 25 °C. Interestingly, in contrast to animals cultured at 15 and 20 °C, the mean bead size slightly decreased after the rehabilitation regime at 25 °C (**Figure 6c**), suggesting that recovery at 25 °C does induce some regenerative effect. The regeneration results observed in animals cultured at 20 °C and recovered at 25 °C (presented in the previous section) support the idea that regeneration at 25 °C is possible, but is likely slower for the population cultured at 25 °C pre cold-shock. Such delayed regeneration could stem from the more drastic difference between the baseline and cold-shock temperature. Finally, these experiments corroborate that cold-shock induced beading occurs in the farthest regions of the neuron, as inter-bead distance increases with cold-shock, and is then reduced after rehabilitation for all culture temperatures (**Figure 6d**). The percentage of small beads (area < 100 pixels) and the percentage of beads with close neighbors (inter-bead distances < 300 pixels) (**Figure S5b**,**d**) also show a reversal of the cold-shock exposure effect in all three physiological temperatures.

Consistent with our previous results, the anterior region of PVD showed a higher number of protrusions than the posterior (**Figure 6e**). Both regions, recapitulate the trends observed for pre cold-shock, post cold-scock, and post rehabilitation in the entire animal. The anterior region consistently exhibits ∼20-100% higher number of beads than the posterior, with pre cold-shocks showing the largest difference. The average bead size does not show differences between these regions (**Figure S5f**). However, similar to previous experiments, the protrusions are more densely distributed in the anterior part, as is expected for a higher number of beads (**Figure 6f**). The results from this assay support our hypothesis that the culture temperature impacts how nematodes respond to acute cold-shock. Animals cultured at 15 °C exhibited the least neurodegenerative signs and faster recovery, while those grown 25 °C showed more drastic beading and slower rehabilitation rate. This difference in response indicates that the magnitude of the cold-shock (based on the baseline temperature) correlates with the induced neurodegenerative effects through a yet unknown mechanism.

### 2.6 Predicting biological status using deep quantitative classification

The quantitative analysis of beading induced by aging and cold-shock indicate that the induced patterns of PVD degeneration are different. To further investigate the morphological changes observed, we took advantage of the rich information obtained from the Mask R-CNN segmentation and feature extraction pipeline, which includes all 46 metrics. Through visual inspection of the raw images, as well as the quantitative analysis of the beading patterns, it is clear that beading phenotypes cannot be fully described with a single feature, such as number of beads. Furthermore, there is significant variability within a population. As shown in **Figure 7a**, a large fraction of aged animals exhibit less than 70 beads, which is considerably lower than the average of the population and is closer to the number of beads for young individuals. Likewise, some young animals showed more than 70 beads, which is significantly higher than the average of the population. The same variability was observed in cold-shock experiments, suggesting that the number of beads does not offer a comprehensive description about biological status of a nematode. Combining two metrics such as number of beads and average bead size still does not provide enough information to distinguish between young and aged adults (**Figure 7a**).

**Figure 7.**
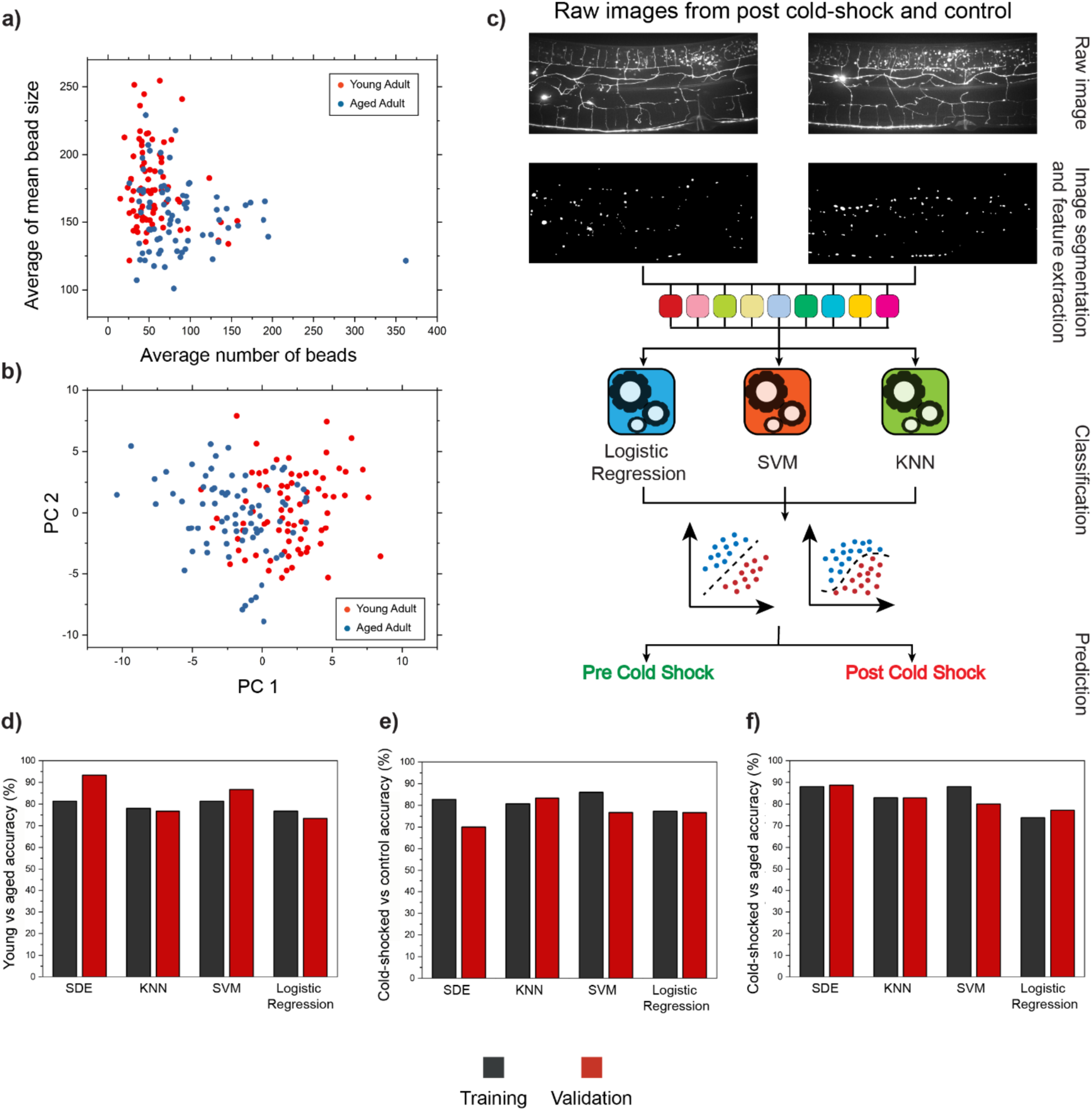
Biological status of a nematode can be predicted based on PVD neuron’s health. **a)** Average of mean bead size versus the average number of beads for young and aged nematodes. Age-induced PVD degeneration patterns are complex and two metrics are not sufficient to accurately classify the two populations. **b)** Principle component analysis (PCA) for young and aged adults does not enable distinguishing young and aged groups, based on the two first principal components. **c)** Schematic of the pipeline for computer-based machine learning models to predict the nematode’s biological status based on the morphological structure of PVD. Raw images are fed to Mask R-CNN algorithm to obtain binary mask, which is then used to extract the 46 metrics. Multiple models were trained based on these 46 metrics, and tested on separate data sets. **d-f)** Classification accuracy for young vs. aged, cold-shocked vs control, and cold-shocked vs. aged nematodes. SDE=Subspace discriminant ensemble, KNN=K-nearest neighbor, SVM=Support vector machine.

Given that beading patterns relay information about the health state of PVD, we reasoned that beading phenotypes could be used to predict the biological state of the animals. To test this hypothesis, we sought to incorporate all 46 metrics extracted from each image in a classification model. In first attempt, as shown in **Figure 7b**, we performed PCA (Principal Component Analysis) on the 46 metrics. Two principal components (PC1 and PC2) explain 46% of the total variance, and are unable to accurately differentiate nematodes from these two stages in their life-span. Thus, we aimed to test the ability of classification models to distinguish young and old nematodes using the metrics extracted from PVD beading patterns. As shown in **Figure 7c**, animals from different groups (e.g. pre and post cold-shock), can exhibit very similar beading patterns. Successful predictive models would prove the presence of subtle neurodegenerative patterns that can only be described using multiple metrics. We first developed a classification model to distinguish young versus old adults. To create a labeled training set, data from the posterior side of PVD for worms younger than 4 days old were grouped together while the second class was comprised of information from nematodes older than 4 days old. An independent validation data set was then generated to test classification accuracy. It should be noted that these two classes are more difficult to distinguish than comparing day 2 vs. day 12 animals (i.e., the youngest vs the oldest samples). We tested four classification algorithms: Subspace Discriminant Ensemble (SDE), Support Vector Machines (SVM), Logistic Regression, and K-Nearest Neighbors (KNN). Two models, SDE and SVM, achieved both training and validation accuracies above 80%, with the validation accuracy of SDE reaching 90% (**Figure 7d**). For age-based classification, the information acquired from the PVD anterior side was also used to train separate models leading to training and validation accuracies higher than 80% (**Figure S6a**). These results suggest that age-induced PVD neurodegeneration causes subtle morphological changes that can only be captured using quantitative deep phenotyping. Similarly, we tested models for classifying nematodes exposed to cold-shock from those that did not experience this stressor. The training and validation set for this analysis was comprised of data from cold-shock performed at all three pre cold-shock temperatures and, as shown in **Figure 7e**, ∼80% classification accuracy was obtained both in training and validation. Since differences between degenerated (i.e., old or cold-shocked) and healthy (young or non cold-shocked) animals have been shown, it was expected that these populations are distinguishable. However, given the significant variability in each population, the high classification accuracy obtained was surprising and points to consistent phenotypic patterns exhibited upon degeneration that are not evident to visual inspection.

To further test the power of our deep phenotyping pipeline, we next investigated potential differences in PVD neurodegeneration exhibited upon aging and acute cold-shock. We compiled data from the anterior and posterior part of the PVD from aging and cold-shock assays to generate training and validation sets. As shown in **Figure 7f**, the SDE model reaches ∼90% training and validation accuracy for the anterior and ∼80% for the posterior regions (**Figure S6c**). This difference in classification accuracy could stem from the anterior part of PVD undergoing stronger beading patterns than the posterior. Notably, these results indicate that these two stressors cause distinct neurodegeneration patterns which can be captured by in depth quantitative analysis. As a last test, we sought to establish whether the differences in bead patterning between the anterior and posterior part of the PVD could be used to classify images of each class. An accuracy of ∼85-90% was achieved from different models, confirming underlying beading pattern differences between these two regions of the neuron (**Figure S6d**). This analysis reveals that at least some of the phenotypic changes in beading patterns are specific to each neurodegenerative stressor. The developed classification models are a powerful tool to identify potential differences in neurodegenerative patterns caused by various environmental stressors (cold-shock or aging).

## 3. Conclusions

Neurons undergo degeneration during the aging process. In *C. elegans*, PVD, a neuron responsible for mechano-sensation and thermo-sensation experiences morphological and functional changes as animals age^32,34^. Prior studies have identified changes in dendrite morphology characterized by disorganization in the menorah-like dendritic arbors^28,29^. In addition, Lezi et al. identified that aging results in the formation of protrusions (or beads) along PVD dendrites^34^. However, analyzing such morphological changes is challenging. Manual inspection to quantify the number of protrusions is time-consuming, labor-intensive, low-throughput, and subject to human bias. In addition, manual counting provides limited information and makes thorough analysis of the complex phenotypes acquired in fluorescence images unfeasible. In order to track morphological changes that PVD undergoes with degeneration, we integrated a cutting-edge deep learning technique to segment the protrusions that form along PVD. This technology decreased the time required to process each image from 3 hours to less than a minute, while eliminating the human bias in analyzing the data. In addition, a secondary algorithm was developed to extract 46 different metrics that make up a comprehensive phenotypic profile that describes the neurodegenerative beading patterns.

We implement a Convolutional Neural Network based algorithm (Mask R-CNN) to carry out challenging image segmentation, unfeasible with traditional image processing approaches. The algorithm segmentation precision and recall achieved 88% and 91% respectively. An important advantage offered by this technology (which cannot be quantified using the metrics above), is its capability to distinguish autofluorescent lipid droplets from actual protrusions, in spite of their remarkable similarities in shape, intensity, and location. The in-depth quantification of PVD morphology enabled by this technology revealed subtle neurodegenerative changes induced by aging and upon exposure to acute cold-shock. With this approach, we identified an increase in the number of beads formed along PVD as animals aged, recapitulating earlier work by Lezi et al^34^. In addition, the reduction in average bead size and inter-bead distance quantified in later points of the nematode’s life-span suggested that the protrusions formed due to aging tend to be small and appear close to each other.

Prior work has focused on the effect of acute cold-shock on a population’s survival and on PVD degeneration independently^32,34,54,66^. However, the neurodegenerative impacts of acute cold-shock on PVD were still unexplored. We sought to test the effect of acute cold-shock on PVD by exposing populations of worms to 4 °C and subsequently quantifying the protrusions generated as a result. We demonstrate that exposure to cold-shock for 16 hrs. or more induces bead formation in PVD. In contrast to the beading patterns induced by aging, the average inter-bead distance increased in animals as a result of cold-shock, a counterintuitive result as an increased bead density is expected with a higher number of beads. This finding, however, can be explained by the formation of beads in the farthest regions of the neuron. These results were the initial signs of aging and cold-shock inducing phenotypically distinct neurodegenerative patterns. We next sought to study the regenerative potential of PVD post cold-shock. Thus, populations of worms exposed to cold-shock were transferred to 3 different temperatures (15, 20, and 25 °C) for a day of recovery. Interestingly, a decrease in the number of beads was observed after the rehabilitation in all 3 temperatures, while the population cultured at 25 °C exhibited the greatest decrease. The increased inter-bead distance induced by cold-shock was reversed in all three temperatures. These results suggest that bead formation due to cold-shock is a reversible process, at least at the earlier stages of adulthood. We also investigated whether culture temperature impacts the severity of bead formation due to cold-shock. Our data suggested that populations cultured at lower temperatures experience less drastic neurodegeneration, while those cultured at higher temperatures undergo more severe damage.

Finally, we use our deep phenotyping approach to predict the biological status of nematodes based on 46 metrics extracted from the images. We tested multiple algorithms (SVM, KNN, SDE, and Logistic Regression) to classify young and old adults, cold-shocked and non-shocked nematodes, and cold-shocked and aged worms. These models achieved ∼85% classification accuracy, indicating distinct beading patterns result from different stressors. Importantly, this classification method, which relies on multiple descriptive metrics of beading patterns, enables deeper exploration of the relevant parameters that describe the biological status of the neuron and its particular degeneration pattern. These promising results suggest that this approach can be used in future studies to characterize beading patterns associated with other conditions or environmental stressors. While the nature of the beads is still unclear, this approach will be crucial in understanding their role, composition, and generation mechanisms, by applying it in genetic or drug screens, and to test the beading patterns formed under other conditions.

In this work, we developed a computer-based comprehensive pipeline to study PVD neurodegeneration in a high-content, automated manner. Our quantitative analysis enabled interrogating the morphological changes that PVD undergoes under different scenarios, leading to deeper understanding of neuronal degeneration. Through this deep phenotyping pipeline, we identify a new environmental stressor (cold-shock) that induces neurodegeneration characterized by beading and reveal distinct neurodegeneration patterns induced by aging. The presented results are evidence that this high-content phenotyping technology can be used to characterize subtle, and noisy degeneration beading patterns with differences amongst stressors unnoticeable to the human eye. This pipeline is a promising approach to further explore the mechanisms underlying of beading neurodegeneration in these and other contexts (such as oxidative stress, dietary restriction, and neurodegenerative disease models), to understand the differences that lead to distinct aging and cold-shock induced degeneration, and to identify whether beads are a result of loss of neuronal integrity or could act as a protective mechanism.

## 4. Materials and Methods

### Worm Culture

The *C. elegans* strain used in this work is NC1686 (wdls51 [F4H12.4::GFP + unc-119(+)]), which expresses GFP in PVD. All populations were cultured on solid Nematode Growth Media (NGM) plates. For aging experiments, 12 mg of Fluorodeoxyuridine (FUdR) was added to 1L of media (50 µM). Animals exposed to this concentration of FUdR produced non-viable eggs. For cold-shock experiments, plates without FUdR were used since experiments took place in 4 days. Age-synchronized populations were obtained by extracting eggs from gravid hermaphrodites using a bleaching solution (1% NaOCl and 0.1 M NaOH). Eggs were then transferred to NGM plates seeded with *E. coli* OP50. M9 buffer (3 g KH_2_PO_4_, 6 g Na_2_HPO_4_, 5 g NaCl, and 1 mL of 1 M MgSO_4_ in 1 L of water) with 5 µM Triton X-100 was used to transfer worms.

### Microscopy

Animals were mounted on 2% agarose pads on glass slide. Agarose pads were placed at room temperature overnight before microscopy. A drop of 10 mM Tetramisole in M9 buffer was added for immobilization. Images were acquired on a Leica DMi8 equipped with a spinning disk confocal head (CrestOptics X-light V2) and a Hamamatsu Orca-Fusion camera using a 63x objective. The illumination source is a Laser Diode Illuminator (89 North LDI). The imaging settings were maintained constant for all images (exposure time of 60ms and laser power at 50%). Due to small field of view provided by the high magnification 63x objective (NA=1.40), two sections of each worm (anterior and posterior of PVD cell body) were imaged separately to cover larger area of the body. Images were acquired as z-stacks of 31 slices taken 1 µm apart. The final raw images used in this study were maximum projections of the z-stacks taken at every one-micron step.

### Image segmentation and analysis

The input to the Mask R-CNN machine learning algorithm trained for this study were 2048×2048 maximum projection PNG images. Images were preprocessed before being fed to the algorithm using MATLAB image processing toolbox (imadjust function) to equalize the image contrast throughout the data set. We modified the Mask R-CNN implementation open-sourced by Matterport Inc. under the MIT license^71^ using Python3, Keras^72^, and Tensorflow^73^. During training, each 2048 x 2048 x 1 image and its set of corresponding binary instance masks were split into 9 overlapping tiles of size 1024 x 1024 x 1. Only non-zero instance masks were kept. The trained Mask R-CNN model was used to predict instance masks by similarly tiling the testing images. Predictions were made sequentially on 9 tiles from the top left to the bottom right of each image, and newly predicted instance masks were kept only if they did not overlap with any previously predicted mask by more than 30%. Only objects yielding a predicted probability greater than 0.9 of being in the foreground or “bead” class were kept. The Mask R-CNN head was trained for 20 epochs and the entire model was trained for 400 epochs, starting from pretrained ImageNet weights. The training data were augmented using random combinations of flips, 90 degree rotations, and affine shearing. The model with the lowest validation loss after 400 epochs was used for predicting instance masks. The binary masks acquired by performing image segmentation using the Mask R-CNN were then coupled with raw images and fed to secondary MATLAB based algorithm to extract metrics describing the morphology of neuron.

### Aging assay

Eggs extracted from gravid hermaphrodites were transferred to a seeded plate and maintained at 20 °C until the population reached late L4 stage and then transferred to an FUdR plate. FUdR plates were checked daily to ensure no viable eggs or progeny were produced. During the first 7-8 days of adulthood nematodes were transferred to a fresh FUdR plate on a daily basis to provide worms with sufficient food specially during their early adulthood. Every 2 days, a subset of nematodes was picked to perform high-resolution microscopy.

### Cold-shock assay

The cold-shock experiments were designed to be conducted in ∼4 days, which included Pre/Post cold-shock microscopy and rehabilitation. For the first two cold-shock assays, eggs extracted from gravid hermaphrodites were transferred to NGM plates and cultured for 4-5 days at 20 °C until they reach day 2 of adulthood, when pre cold-shock microscopy was performed. Cold-shock was performed by transferring plates to a 4 °C refrigerator for the designated amount of time. Plates were then placed at room temperature for 1 hr. before performing post cold-shock microscopy. This hour-long rehabilitation allowed nematodes to regain their mobility. For the tests where rehabilitation was needed, plates were transferred to designated temperature (15 °C, 20 °C, and 25 °C) for one day before post-rehabilitation microscopy was performed. For the pre cold-shock culture temperature effect assay, nematodes were cultured at 20 °C until reaching young adulthood. Subsequently, the three populations were transferred to 15°C, 20 °C, and 25°C incubator and cultured for 3.5, 2.5, and 1.5 days before performing pre cold-shock microscopy on each population (to ensure all three samples reach the same developmental stage). Populations experienced 16 hrs. of cold-shock at 4 °C prior to post cold-shock microscopy. The samples were then transferred back to the temperature they were cultured at before cold-shock for one-day to examine the post shock recovery.

### Principal Component Analysis and classification

Principal Component Analysis (PCA) based on correlation was performed using JMP Pro 14 software. For this analysis, a data set comprised of 150 images (half from animals younger than 4 days and half from animals older than 4 days old). All 46 metrics extracted from images were incorporated in the analysis. The first two principal components explained 46% of the variance. Classification of biological status was conducted using MATLAB Classification Learner App. For all training sessions, all 46 metrics extracted from images were incorporated to train the models, and used 5 folds cross-validation was carried out. A separate validation set was used to test performance.

## 6. Acknowledgements

This work was supported by the National Institutes of Health (R00AG046911 and R21AG059099) and the National Science Foundation (IOS1838314). The *C. elegans* strain was provided by the *Caenorhabditis* Genetics Center, which is funded by NIH Office of Research Infrastructure Programs (P40 OD010440). We thank Dong Yan and E Lezi for useful discussions.

## 7. Contribution

S.S. and A.S.M. conceived and designed the study. S.S. conducted the experiments. K.B.F. trained the machine learning algorithm. S.S., A.S.M., and K.B.F. analyzed the data and wrote the manuscript.

