## Supplementary Information for "Deep learning-enabled phenotyping reveals distinct patterns of neurodegeneration induced by aging and cold-shock"

**46 metrics extracted from each image:**

1. Number of beads
2. Total area occupied by beads
3. Average bead size
4. Standard deviation of bead size
5. Standard deviation of mean of bead size (Standard deviation/Mean)
6. Standard error of mean for bead size
7. 90th percentile of bead size
8. 75th percentile of bead size
9. 50th percentile of bead size
10. 25th percentile of bead size
11. Average bead size for beads larger than 100 pixels
12. Average bead size for beads smaller than 100 pixels
13. Number of beads with area larger than 100 pixels
14. Number of beads with area smaller than 100 pixels
15. Percentage of beads with area larger than 100 pixels
16. Percentage of beads with area smaller than 100 pixels
17. Average inter-bead distance
18. Standard deviation of inter-bead distance
19. Standard deviation of mean inter-bead distance (Standard deviation/Mean)
20. Standard error of mean for inter-bead distance
21. Percentage of beads with inter-bead distance less than 150 pixels
22. Percentage of beads with inter-bead distance less than 300 and greater than 150 pixels
23. Percentage of beads with inter-bead distance less than 450 and greater than 300 pixels
24. Percentage of beads with inter-bead distance greater than 450 pixels
25. Median of bead size
26. Maximum of bead size

27. Average size for beads larger than 100 pixels/ Average size for beads smaller than 100 pixels
28. Mean size for smallest beads (Smallest beads are smaller half of the beads)
29. Mean size for largest beads (Largest beads are larger half of the beads)
30. Mean size of largest beads/ Mean size of smallest beads
31. Average of mean bead intensity
32. Median of mean bead intensity
33. Max of mean bead intensity
34. Standard deviation of mean bead intensity
35. Standard deviation of mean bead intensity (Standard deviation/Mean)
36. Standard error of mean of bead intensity
37. $90_{th}$ percentile of mean bead intensity
38. $75_{th}$ percentile of mean bead intensity
39. $50_{th}$ percentile of mean bead intensity
40. $25_{th}$ percentile of mean bead intensity
41. Mean intensity of smallest beads
42. Mean intensity of largest beads
43. $90_{th}$ percentile of inter-bead distance
44. $75_{th}$ percentile of inter-bead distance
45. $50_{th}$ percentile of inter-bead distance
46. $25_{th}$ percentile of inter-bead distance

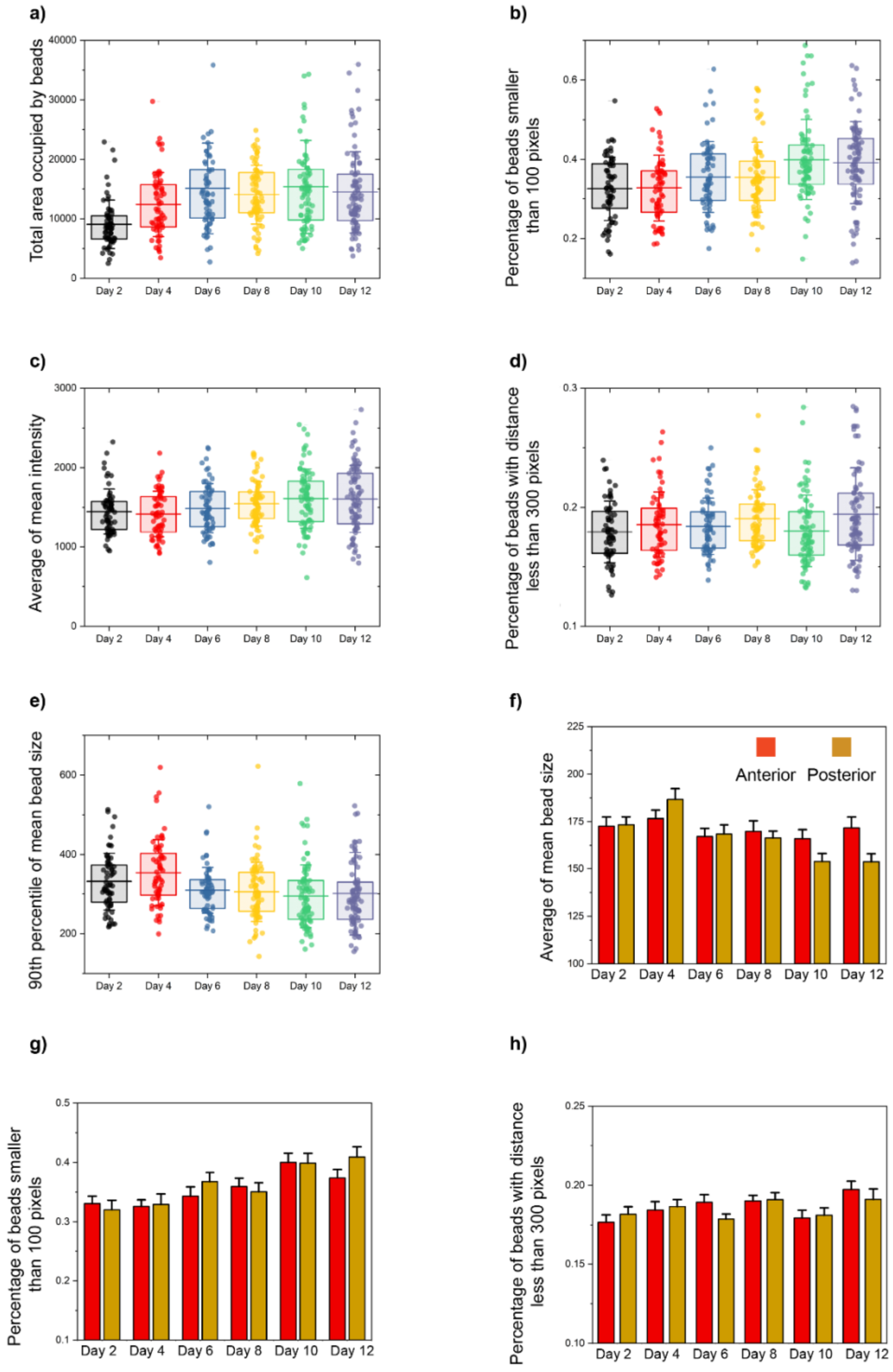


***Figure 1.*** *Age-induced degeneration causes morphological variation on PVD structure.* ***a)*** *Total area of PVD neuron covered with beads.* ***b)*** *Percentage of beads with size lower than 100 pixels.* ***c)*** *Average of the mean intensity of the beads.* ***d)*** *The percentage of inter-bead distances lower than 300 pixels.* ***e)*** *90th percentile of beads fluorescence intensity. Lines are 25th percentile, mean, and 75th percentile. Whisker is the standard deviation.* ***f-h)*** *Average of mean bead size, percentage of beads with inter-bead distance less than 300 pixels, and percentage of beads with size lower than 100 pixels for anterior versus posterior part of the PVD neuron.*

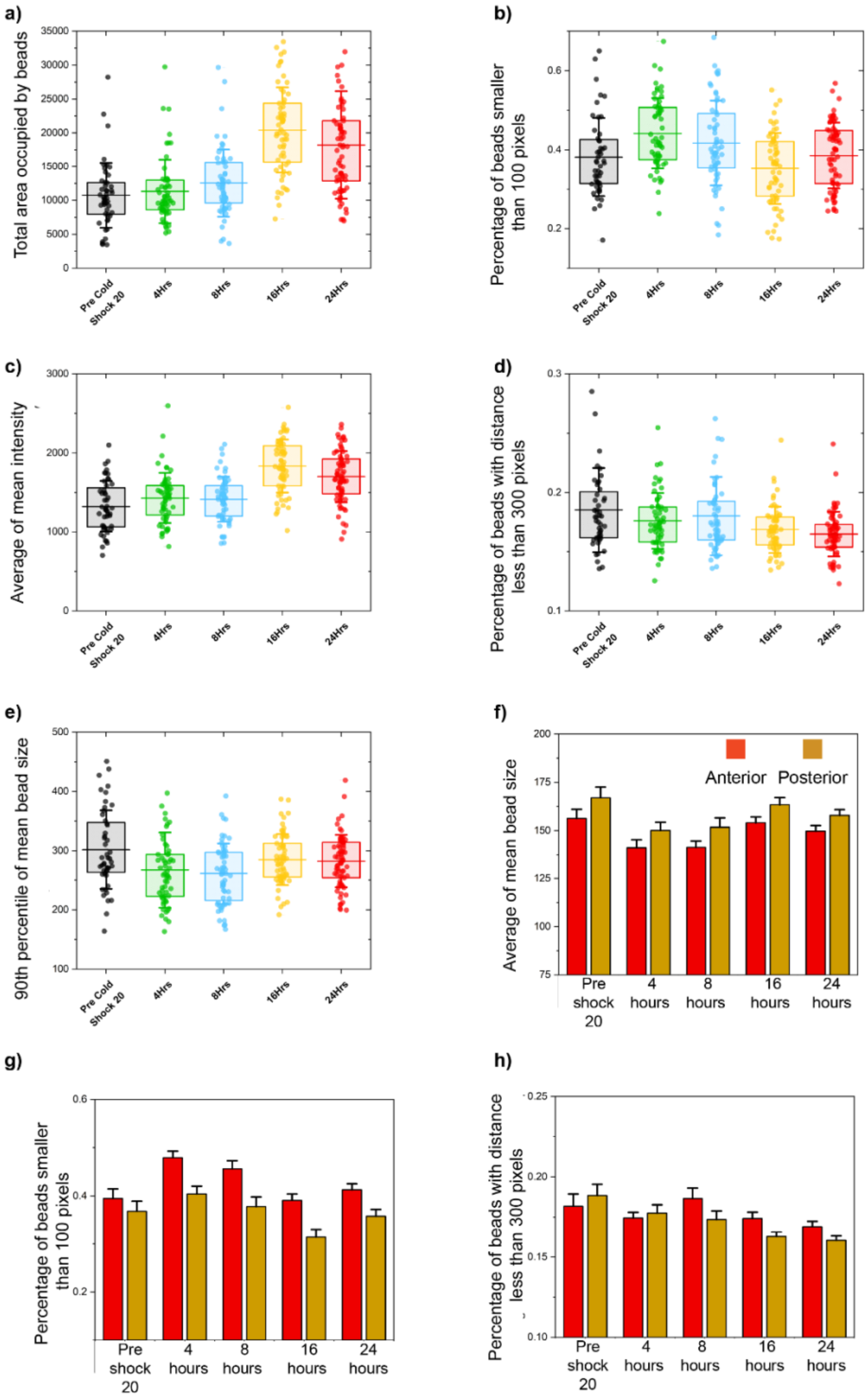


***Figure 2.*** *PVD neuronal structure undergoes morphological changes upon exposure to acute cold-shock.* ***a)*** *Total area of PVD neuron covered with beads.* ***b)*** *Percentage of beads with size lower than 100 pixels.* ***c)*** *Average of the mean intensity of the beads. This line is a transcriptional line; thus, the change is the representative of the fluctuation in concentration on promoter along beads.* ***d)*** *The percentage of inter-bead distances lower than 300 pixels.* ***e)*** *90th percentile of beads fluorescence intensity. The lines are 25th percentile, mean, and 75th percentile. Whisker is standard deviation.* ***f-h)*** *Average of mean bead size, percentage of beads with inter-bead distance less than 300 pixels, and percentage of beads with size lower than 100 pixels for anterior versus posterior part of the PVD neuron.*

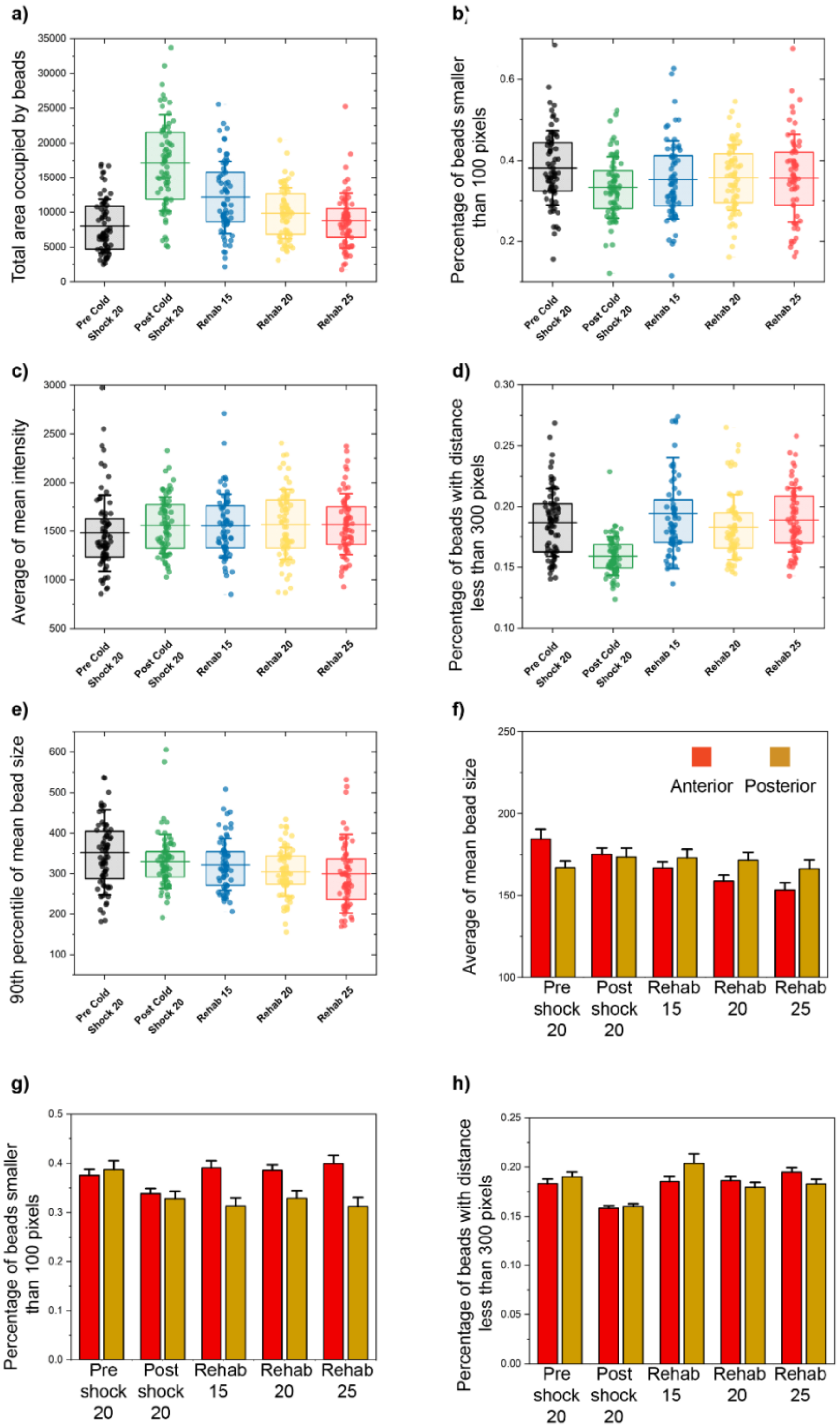


***Figure 3.*** *PVD neurodegenerative phenotypes caused by acute cold-shock is reversible and can be alleviated by post shock rehabilitation.* ***a)*** *Total area of PVD neuron covered with beads.* ***b)*** *Percentage of beads with size lower than 100 pixels.* ***c)*** *Average of the mean intensity of the beads. This line is a transcriptional line; thus, the change is the representative of the fluctuation in concentration on promoter along beads.* ***d)*** *The percentage of inter-bead distances lower than 300 pixels.* ***e)*** *90th percentile of beads fluorescence intensity. The lines are 25th percentile, mean, and 75th percentile. Whisker is the standard deviation.* ***f-h)*** *Average of mean bead size, percentage of beads with inter-bead distance less than 300 pixels, and percentage of beads with size lower than 100 pixels for anterior versus posterior part of the PVD neuron.*

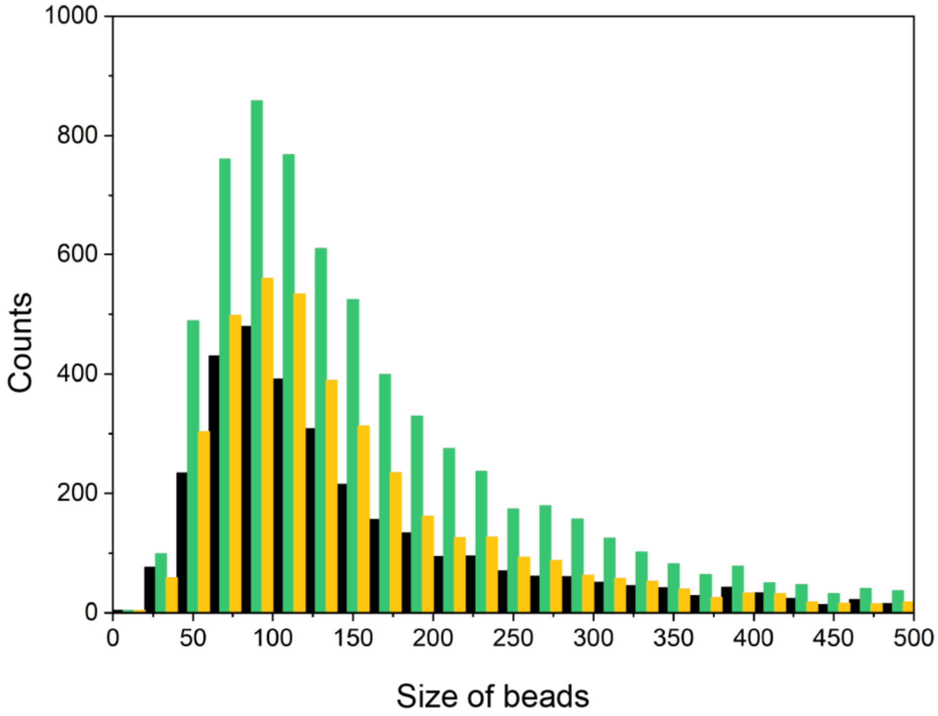


***Figure 4.*** *Histogram distribution of individual bead size for recovery assay of a population cultured at 20 °C.*

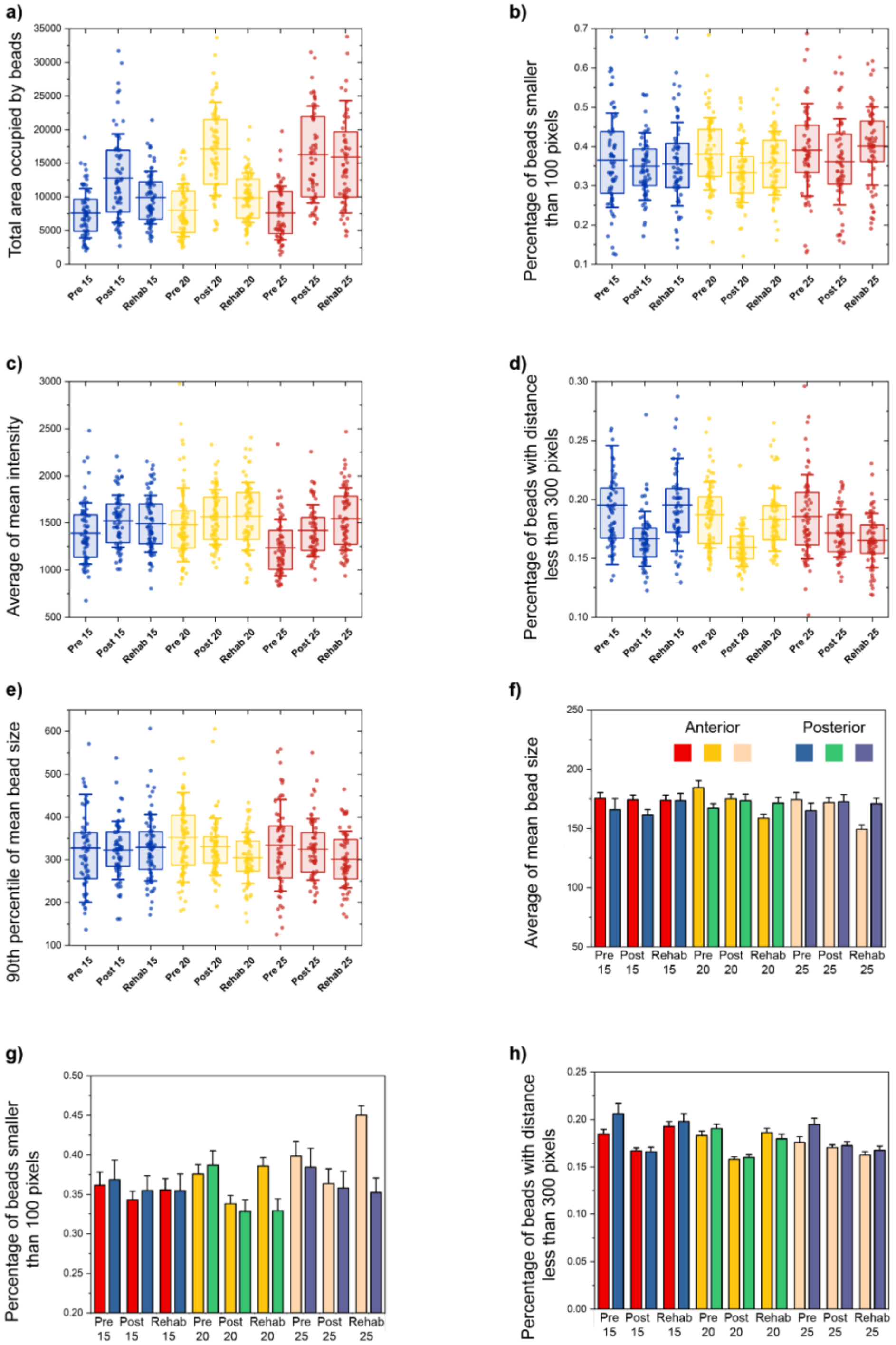


***Figure 5.*** *Populations cultured at different temperature before being exposed to cold-shock respond in various ways to this external stressor.* ***a)*** *Total area of PVD neuron covered with beads.* ***b)*** *Percentage of beads with size lower than 100 pixels.* ***c)*** *Average of the mean intensity of the beads. This line is a transcriptional line; thus, the change is the representative of the fluctuation in concentration on promoter along beads.* ***d)*** *The percentage of inter-bead distances lower than 300 pixels.* ***e)*** *90th percentile of beads fluorescence intensity. The lines are 25th percentile, mean, and 75th percentile. Whisker is standard deviation.* ***f-h)*** *Average of mean bead size, percentage of beads with inter-bead distance less than 300 pixels, and percentage of beads with size lower than 100 pixels for anterior versus posterior part of the PVD neuron.*

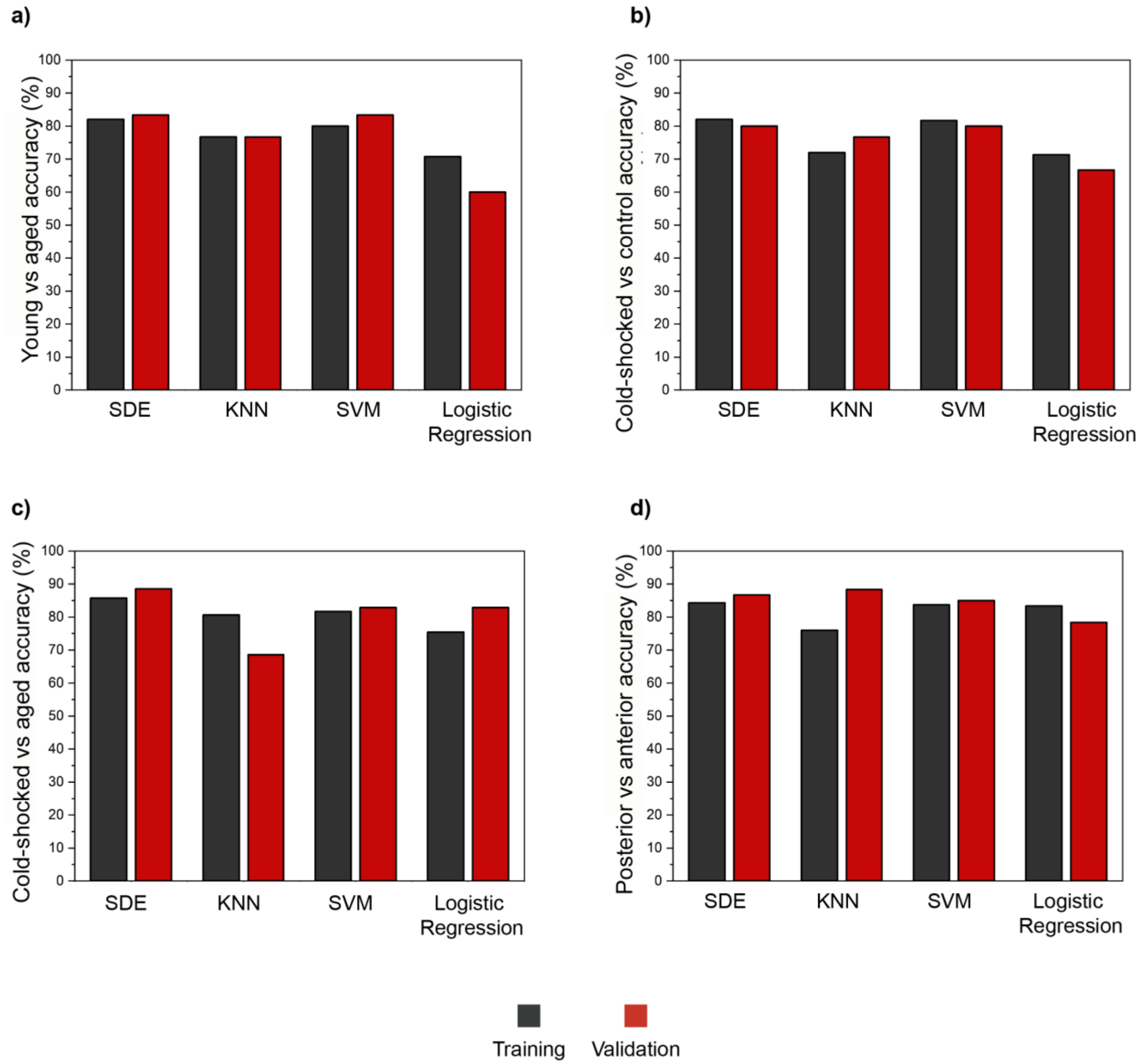


***Figure 6.*** *Biological status of a nematode can be predicted based on PVD neuron's health.* ***a)*** *Classification accuracy for young vs aged nematodes. The images were acquired from anterior part.* ***b)*** *Classification accuracy for cold-shocked vs control nematodes. The images were acquired from posterior part.* ***c)*** *Classification accuracy for cold-shocked vs aged nematodes. The images were acquired from posterior part.* ***d)*** *Classification accuracy for posterior vs anterior images of nematodes.*
